# Do mechanical strain magnitude and rate drive bone adaptation in adult women? A 12-month prospective randomized trial

**DOI:** 10.1101/421156

**Authors:** Karen L. Troy, Megan E. Mancuso, Joshua E. Johnson, Zheyang Wu, Thomas J. Schnitzer, Tiffiny A. Butler

**Affiliations:** Department of Biomedical Engineering, Worcester Polytechnic Institute, Worcester, MA; Orthopaedic Biomechanics Research Laboratory, University of Iowa, Iowa City, IA; Department of Mathematical Sciences, Worcester Polytechnic Institute, Worcester, MA; Department of Physical Medicine and Rehabilitation, Northwestern University, Chicago, IL

**Keywords:** Bone QCT/μCT, Bone modeling and remodeling, Exercise, Biomechanics, Clinical Trials

## Abstract

Although there is strong evidence that certain activities can increase bone density and structure in some individuals, it is unclear what specific mechanical factors govern the response. This is important because understanding the effect of mechanical signals on bone could contribute to more effective osteoporosis prevention methods and efficient clinical trial design. The degree to which strain rate and magnitude govern bone adaptation in humans has never been prospectively tested. Here, we studied the effects of a voluntary upper extremity compressive loading task in healthy adult women during a twelve month prospective period. One hundred and two women age 21-40 participated in one of two experiments. (1): low (n=21) and high (n=24) strain magnitude. (2): low (n=21) and high (n=20) strain rate. Control: (n=16): no intervention. Strains were assigned using subject-specific finite element models. Load cycles were recorded digitally. The primary outcome was change in ultradistal integral bone mineral content (iBMC), assessed with QCT. Interim timepoints and secondary outcomes were assessed with high resolution pQCT (HRpQCT). Sixty-six subjects completed the intervention, and interim data were analyzed for 77 subjects. Both the low and high strain rate groups had significant 12-month increases to ultradistal iBMC (change in control: -1.3±2.7%, low strain rate: 2.7±2.1%, high strain rate: 3.4±2.2%), total iBMC, and other measures. “Loading dose” was positively related to 12-month change in ultradistal iBMC, and interim changes to total BMD, cortical thickness and inner trabecular BMD. Subjects who gained the most bone completed, on average, 130 loading bouts of (mean strain) 550 με at 1805 με/s. Those with the greatest gains had the highest loading dose. We conclude that signals related to strain magnitude, rate, and number of loading bouts contribute to bone adaptation in healthy adult women, but only explain a small amount of variance in bone changes.

## Introduction

Exercise-based interventions have long been considered a viable option for preserving and enhancing bone strength ^(1)^ because bone adapts to best resist its habitual mechanical loading environment. Individuals who play sports and load with odd strains (soccer, squash) have been observed to have better bone mechanical properties than those who do not ^(2)^. Furthermore, clinical trials have shown that high impact activities such as jumping and hopping can improve bone density in growing children ^(3)^ and young adults ^(4)^ and maintain bone density in older adults ^(5)^. However, while the evidence is strong that certain activities can increase bone density and structure in some individuals, it is not clear what specific mechanical factors govern the response. Furthermore, these factors interact with each individual person’s physiology to create a variable response, which is not well understood.

Animal *in vivo* loading models have demonstrated that mechanical signals related to strain rate ^(6–8)^ and strain magnitude ^(9, 10)^ regulate bone adaptation. There is no consensus on which specific signal(s) osteocytes sense; evidence supports lacunar-cannilicular fluid flow ^(11, 12)^, flow of ions and the resulting electromagnetic signal ^(13)^, direct damage of osteocytes ^(14)^, microdamage of the surrounding bone that results in altered stress or strain ^(15, 16)^ and other candidates ^(17)^. Regardless of the exact mechanism, all of these signals are closely related to (and driven by) mechanical strain. *In vivo* loading models have also established that, to elicit an adaptive response, the mechanical signal must be both dynamic and novel ^(18)^. Despite extensive animal literature, the degree to which mechanical strain magnitude and rate govern bone adaptation in humans has never been prospectively tested.

One major challenge is that bone strain is difficult to measure noninvasively. As a result, indirect measures, such as surveys for physical activity, which include weighting factors based on experimentally measured ground reaction force (GRF) and rate of GRF, have been proposed ^(19, 20)^. Others have proposed “bone loading” indices that are based on similar measures (e.g. accelerometry) ^(21, 22)^. While these can be helpful in identifying the types of activities that should theoretically elicit an osteogenic response, they do not account for individual differences in bone structure, which have a large influence on bone strain ^(23)^. Alternatively, validated subject-specific FE models can provide accurate estimates of bone strain^(24–26)^ when the proper boundary conditions (magnitude, direction, and locations of application) are known.

Our previously validated upper extremity loading model ^(26)^ provides a well-controlled framework to understand the degree to which strain magnitude and rate influence bone adaptation in people. In this model, an individual produces a compressive force through the radius by leaning onto the palm of the hand to achieve a target force. Feedback is given using a scale or loadcell, and individuals are given sound cues to assist in achieving a regular and consistent load/unload cycle. In a pilot group of 19 young adult women, we found that a mean energy equivalent strain of 734 ± 238 με applied 50 cycles per day, 3 days per week elicited modest increases in distal radius bone mineral content (BMC) and prevented seasonal loss of BMC observed in a control group ^(26)^. We also observed that high strain regions of the radius gained significantly more bone than low strain regions, suggesting that the local mechanical signals were, in part, driving the response ^(27)^. Although these results were promising, the study was limited in scope and duration.

Here, our purpose was to quantify the degree to which bone strain influences bone adaptation in the upper extremity of healthy adult women during a twelve month prospective study period. We hypothesized that (1) bone accrual would be proportional to strain magnitude and strain rate, and (2) structural changes would include increased cross-sectional area and cortical thickness, and increased trabecular bone mass near the endosteal surface.

## Methods

### Participant Characteristics

One hundred and two women, age: 28 ± 6 years, height: 164 ± 8 cm, mass: 65 ± 9 kg, were recruited from the community for this mechanistic randomized prospective trial. Healthy women age 21-40 were included in the present study; this group is at peak bone mass ^(28, 29)^, and compared to men, have increased risk of osteoporosis later in life. After initial telephone screening for inclusion criteria, potential subjects were screened with dual energy x-ray absorptiometry (DXA) of the non-dominant radius and circulating levels of 25-hydroxyvitamin vitamin D and estradiol. Exclusion criteria included BMI outside [18-25 kg/m^2^], irregular menstrual cycles, no regular calcium intake, use of medications affecting bone health, history of radius fracture or injury to the non-dominant shoulder or elbow, regular participation (>2 times per month) in activities with high loads at the forearm (e.g. gymnastics, volleyball), 25- hydroxyvitamin D serum levels below 20 ng/ml, and DXA T-score outside [-2.5 to 1.0]. In total, 102 potential participants provided written, informed consent to participate in this institutionally approved study. All subjects were recruited at a single site between December 2013 and June 2017. The trial was conducted in accordance with Good Clinical Practice Guidelines. Compliance and adverse events data were reviewed annually with a study monitor.

### Study Design

This was a 12 month, prospective, mechanistic randomized trial that utilized a distal radius compressive loading intervention to investigate the effect of strain on bone adaptation. After obtaining informed consent, subjects were assigned into either control or one of two exercise arms that manipulated strain magnitude (Experiment 1: low and high strain magnitude) or strain rate (Experiment 2: low and high strain rate, detailed in Table 1). Group assignments were made by drawing slips of paper from an envelope (e.g. LOW, HIGH, CONTROL), and control subjects were randomized during Experiment 1. Exercise groups were instructed to apply 100 cycles of axial force (one bout), four times weekly, by leaning onto the palm of the hand. Loading was accomplished using a custom device, consisting of a uniaxial load cell (Standard Load Cells; Gujarat, India), data logger (DATAQ DI-710), and LED indicators that lit up when the applied force was within ±10 N of the target value. To allow subjects to get used to the intervention, those in the exercise groups were assigned a nominal 200 N target force magnitude for the first three months of loading. Thereafter, a subject-specific target force was prescribed to achieve target strain parameters based on computed-tomography based finite element (FE) models (described below; Figure 1).

**Figure 1.**
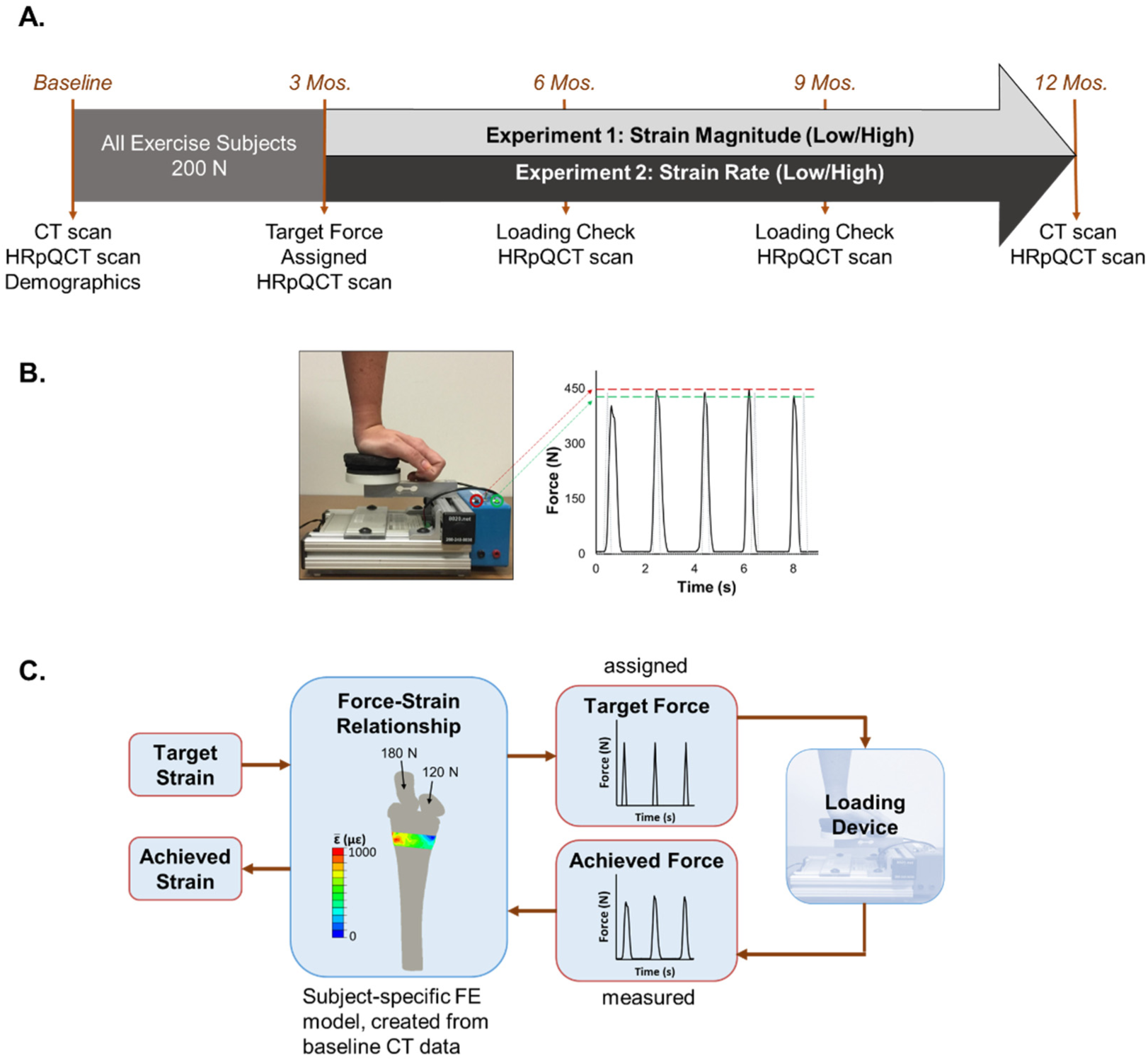
**A**. Summary of the data collection timeline for participants assigned to exercise groups; **B.** Loading device used to manipulate applied force magnitude via feedback lights (green set to target force minus 10 N, red to target force plus 10 N). Loading frequency was controlled using pre-recorded auditory cues. The force vs. time curve shows a representative load cell signal (black) versus ideal assigned loading stimulus (gray), with dashed lines indicating the forces at which feedback is given; **C.** Linear FE model used to estimate energy equivalent strain in the transverse section matching the imaged site. The force-strain relationship was used to assign each subject a target force and calculate the resulting strain from load cell recordings.

**Table 1.**
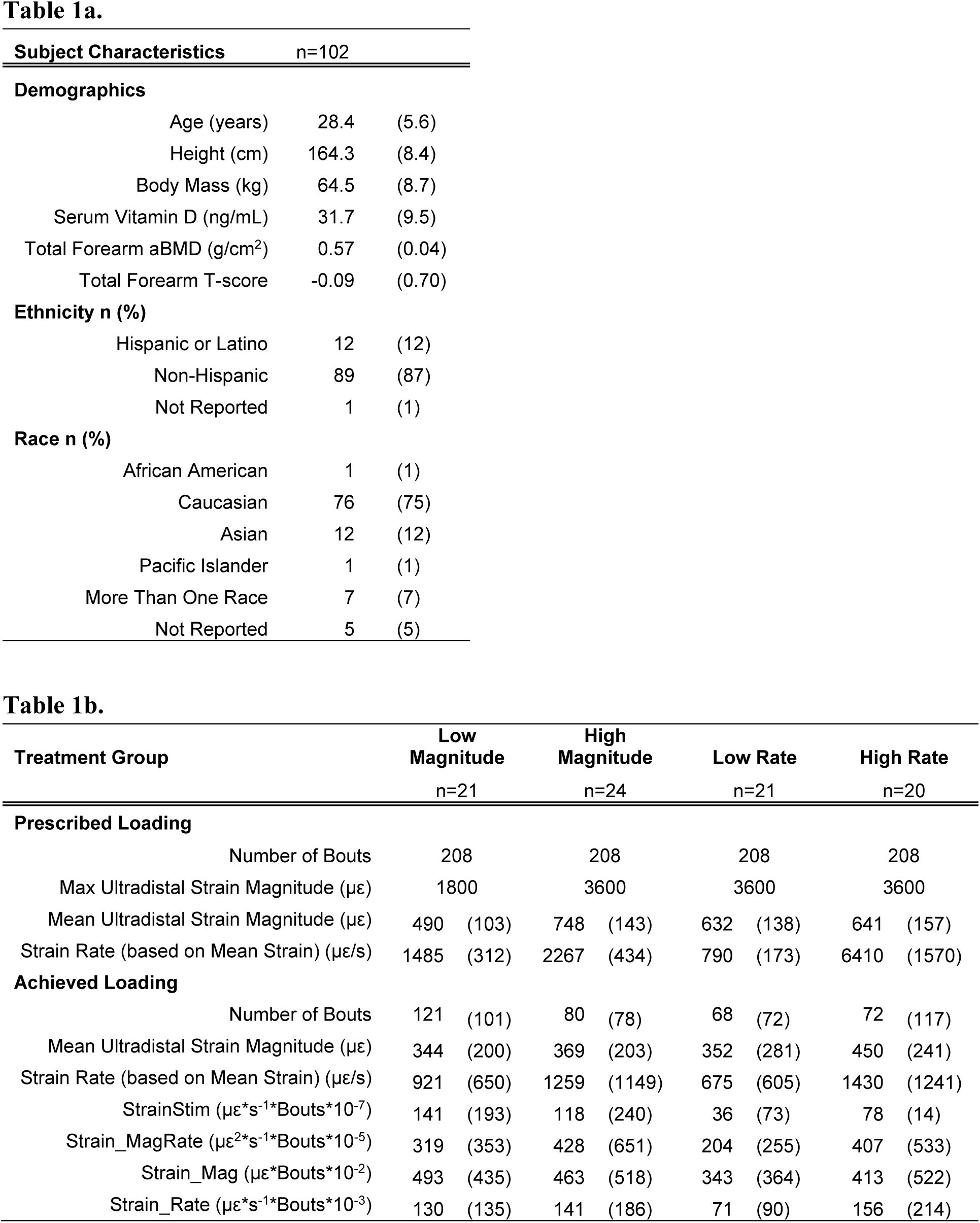
Mean (SD) baseline subject characteristics (a), and loading intervention by group (b). Strain magnitudes were assigned based on the maximum energy-equivalent strain within the ultradistal region, as calculated by FE model. However, for loading dose calculations, the achieved mean energy-equivalent strain was used, since it better represents the strain experienced within the region.

Due to considerations of subject safety, no subject was assigned a force larger than 450 N or what she could comfortably and consistently apply, even if the force required to achieve the target strain was larger than that. Partway through the study, in response to reports of wrist soreness from some subjects, this upper limit was reduced to 350 N. Loading rate and cycle period were controlled using verbal instructions (e.g. “load slowly and evenly” versus “load as rapidly as possible”) and sound cues recorded on a portable voice recorder. Sound cues consisted of 100 beeps (long beeps for the slow rate group, short beeps for the fast rate group) occurring at 2-second intervals. Compliance was monitored every three months using data logger recordings and log books maintained by subjects.

The primary outcome was 12 month change in integral ultradistal radius bone mineral content and bone mineral density (iBMC and iBMD), as measured by quantitative CT analysis (QCT). Secondary outcomes included 12 month changes in other regions, and microstructural measures with interim timepoints. A power analysis based on pilot data ^(26)^ determined that 20 subjects per group would have 80% power to detect a 12 month change in BMC of 1.0±1.1%. Because this study was designed to test a mechanistic outcome and some subjects assigned to loading groups did not engage in any loading at all (making them effectively members of the control group), randomization was adjusted part-way through to reduce the number of subjects assigned to the control group (from 20 to 16), and to increase the size of the loading groups.

### Data Collection

Demographic information and imaging data (DXA, computed tomography; CT, and high resolution peripheral computed tomography; HRpQCT) were collected at baseline. Hand dominance was determined using the Edinburgh inventory ^(30)^ and expressed as left or right decile. CT was collected at baseline and 12 months. HRpQCT was updated every three months during the study period.

### High Resolution Peripheral Quantitative Computed Tomography

Changes in radius microstructure were assessed using HRpQCT (Xtreme CT I, Scanco Medical; Brüttisellen, Switzerland). Bilateral scans were acquired in a standard 9.02 mm region consisting of 110 transverse slices (82 µm isotropic voxel size) beginning 9.5 mm proximal to the distal endplate. Structural changes were measured for the mutually overlapping region, using the manufacturer’s 2D region-matching algorithm. Total mean cross-sectional area (CSA; mm^2^) and total volumetric bone mineral density (Tt.BMD; mgHA/cm^3^) were measured. Trabecular number (Tb.N; mm^−1^), thickness (Tb.Th; mm) and BMD (Tb.BMD; mgHA/cm^3^) were measured using the manufacturer’s standard analysis protocol. The trabecular region was further divided into inner (central 60%; Tb.BMDinn) and outer regions (outer 40%; Tb.BMDmeta). In our lab, the coefficient of variation (CV) for densitometric variables is < 0.3%. Cortical vBMD (Ct.BMD; mgHA/cm^3^), cortical thickness (Ct.Th; mm), and cortical porosity (Ct.Po; %) were calculated using the dual-threshold method ^(31–33)^. The CVs of these variables range from 0.4- 13%. All HRpQCT analyses were blinded to group assignment.

### Quantitative Computed Tomography Analysis

At baseline and 12 months, CT scans of the distal-most 12 cm of the each forearm were acquired (GE Brightspeed, GE Medical, Milwaukee, WI, 120 kV, 180 mA, voxel size 234 µm x 234 µm x 625 µm). A calibration phantom (QRM, Moehrendorf, Germany) with known calcium hydroxyapatite equivalent concentrations was included in the field of view to relate CT attenuation (Hounsfield Units) to equivalent bone density (g/cm^3^).

Changes in bone macrostructure were quantified from CT data using Mimics v15.1 (Materialise, Leuven, Belgium). Follow-up scans were registered to baseline using rigid image registration and the periosteal surface was defined using a 0.175 g/cm^3^ density threshold ^(26)^. Based on methods previously established ^(34)^, we defined integral and endocortical compartments (denoted in QCT variable names with prefixes *i* and *ec*). Briefly, the integral compartment consisted of all voxels within the periosteal surface. The endocortical compartment was comprised of the entire set voxels located within 2.5 mm of the periosteal surface (including all cortical bone). For each compartment bone volume (BV; cm^3^), bone mineral content (BMC; g) and volumetric bone mineral density (BMD; g/cm^3^) were calculated. QCT parameters for the trabecular compartment were not analyzed. Instead, HRpQCT data were analyzed, which provided a greater level of detail. Using previously established methods ^(26)^, we also calculated compressive strength index (CSI; g^2^/cm^4^), and bending strength index (BSI, cm^3^). All parameters were calculated for total and ultradistal regions except for strength measures, which were only calculated for the ultradistal region. The total region extended 45 mm proximal from the subchondral plate and distally to the styloid tip; the ultradistal region extended 9.375 mm proximal from the subchondral plate. The coefficient of variation for these QCT measures in our lab ranges from 0.7 to 2.3% (ultradistal region); 0.3 to 0.6% (total region); 0.9 to 2.3% (strength indices). All QCT analyses were blinded to group assignment.

### Continuum Finite Element Modeling

Finite element models were constructed from the QCT scans using methods validated using cadaveric mechanical testing ^(35)^. Models were used to simulate one cycle of axial loading to determine the subject-specific force needed to achieve the desired target strain within the distal radius. As with our previous work, we used energy-equivalent strain as the measure of interest, since it provides a scalar value that has been related to bone adaptation ^(27, 36)^. Strain values were assigned to each subject based on the maximum energy-equivalent strain within the ultra-distal region of the radius, as calculated using the continuum FE model for that subject. The baseline FE model for each subject was used to adjust the custom loading device so that the LEDs would light up when that individual achieved her target strain. At all subsequent time points, data recorded from the load cell were applied to the FE model to calculate the *actual* mean strain within the region achieved by the subject, based on applied force (Figure 1c).

### Load Cell Analysis

At each follow-up visit, load cell recordings were analyzed using custom code in Matlab. The beginning, peak and end of each loading waveform were identified using a custom algorithm, and the resulting frequency spectrum calculated using a Fast Fourier Transform. Based on subject-specific FE models, frequency data were used to calculate the loading stimulus using the relationship suggested by Turner ^(18)^.

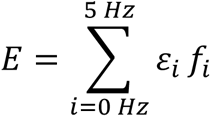

Where *E* is the strain stimulus for the entire loading bout, *f_i_*is the frequency value for bin *i*, and *E_i_* is the peak-to-peak strain magnitude of frequency component *i.* A cutoff of 5 Hz was selected, based on analysis of the load cell frequency content, which showed that over 95% of the signal power was <2 Hz. We also calculated peak-to-peak strain magnitude and strain rate for the loading portion of each cycle for each subject and each loading bout. Because voluntary loading produced variable and sometimes inconsistent loading signals, we evaluated several candidate measures of “loading dose”, which was intended to serve as an overall metric of mechanical loading, considering strain parameters and protocol compliance. We considered the following candidate measures of “loading dose”.

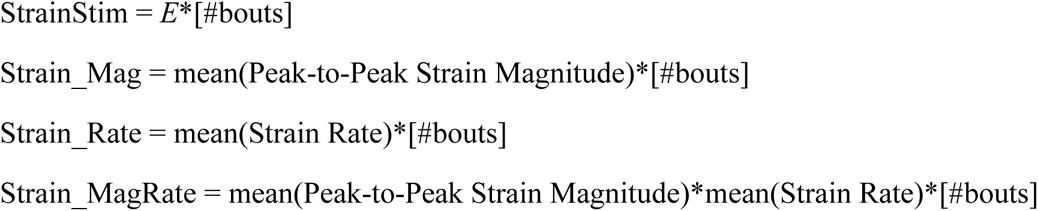

### Statistical Analysis

Descriptive statistics were calculated and data normality was assessed. Group demographics and loading dose received were compared using ANOVA and the Bonferroni-corrected *post hoc* t-test. Analyses were initially by intention to treat. The hypothesis that bone mass would increase proportionally to the applied strain magnitude (Experiment 1) was tested in two ways. First, subjects were analyzed by group (control vs. low and high strain magnitude groups). For the group analysis, the 12-month change in ultradistal iBMC was analyzed as the primary dependent variable in a linear regression model with coefficients representing contrasts between each of the two experimental groups and the control group. The secondary outcome measures were also compared between groups using regression models based on the change scores at each of the time points (change from baseline). Similar analyses were performed to examine the effect of strain rate on bone (Experiment 2).

In the second analysis, we considered “loading dose” achieved by each subject as a continuous variable, with the dose for control subjects being zero. Because dose includes both magnitude and frequency components, all groups were combined into a single regression model with the 12-month change in radius ultradistal iBMC as the primary outcome. The secondary outcome measures were also considered. To test the hypothesis that bone structural changes would include increased cross-sectional area and cortical thickness, and increased endocortical density, these factors were treated as dependent variables in linear regression models, similar to the previous analyses. We assessed the F-statistic of the overall regression, and the t-statistic of each explanatory variable, considering α=0.05 to be significant. As an exploratory *post hoc* analysis, subjects were grouped into tertile based on the change in ultradistal iBMC. Subject demographics, baseline values, and loading dose were compared between tertiles, in order to gain insight into what factors were associated with the most gains in ultradistal iBMC. Bonferroni-adjusted *post hoc* t-tests were used to compare individual tertiles.

## Results

### Participant Characteristics

Baseline characteristics are summarized in Table 1 and were not different between experimental groups. Sixty-six subjects completed the study and were included in the 12-month analysis. Seventy-seven subjects had some follow-up data available and were included in our analyses of interim time points (Figure 2). On average, subjects assigned to one of the loading groups completed 85 (SD: 92) loading bouts in total. However, the total number of loading bouts varied considerably, from 0 to 357. All measures of loading dose were significantly greater for loading groups than for controls (p≤0.046; Table 1)

**Figure 2.**
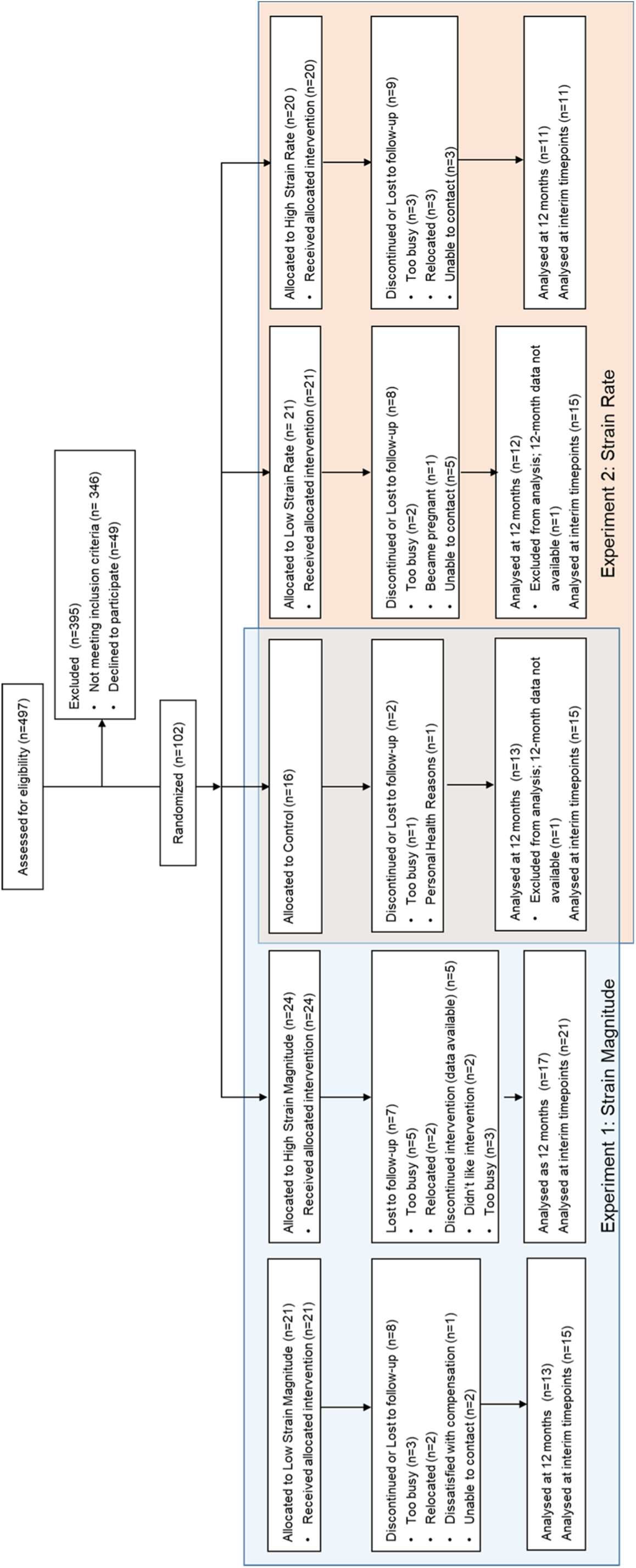
Consort chart describing participant flow.

### Adverse Events

There were no serious adverse events. Temporary soreness of the loaded wrist was the most commonly reported adverse event (28% of subjects; 29 reports). Two of these subjects noted that this briefly affected their daily activities (did fewer chores or avoided exercises that weighted the hands), and one took ibuprofen. Eight subjects reported soreness at other sites (elbow, shoulder, hand), which included aggravation of previous injuries (e.g. shoulder pain from an injury that was several years old) that they thought might be due to the loading intervention. All subjects reported that soreness resolved within 3-14 days. Five subjects reported that pain from previous injuries temporarily prevented them from completing the assigned loading, but did not believe this was caused or aggravated by the intervention. Radiology reports indicated no visible changes in wrist anatomy between initial and 12-month visits. Lack of time or relocation were the most common reasons expressed for dropping out (22 subjects).

### Effect of Strain on 12-month Change in Bone Mass and Structure (QCT)

None of the regression models that included strain magnitude groups were significant for overall model fit, although the membership in the low strain magnitude group was associated with slight gains in ultradistal iBMC (p=0.041), and consistent and significant increases in CSA, iBV and ecBV that indicated periosteal expansion (Table 2). Strain rate had a stronger effect on 12-month change in QCT variables than did strain magnitude (Figure 3). In models comparing the low and high strain rate groups to the control group, both loading groups were significantly and positively associated with the increases to total and ultradistal iBMC, iBMD, ecBMC, and ecBMD. Fifty-six and 52% of the variance in change to ultradistal and total iBMD, respectively, was explained by group membership of these subjects (Table 3). Increases to ultradistal compressive and bending strength indices were significantly and positively associated with Experiment 2 loading group membership.

**Figure 3.**
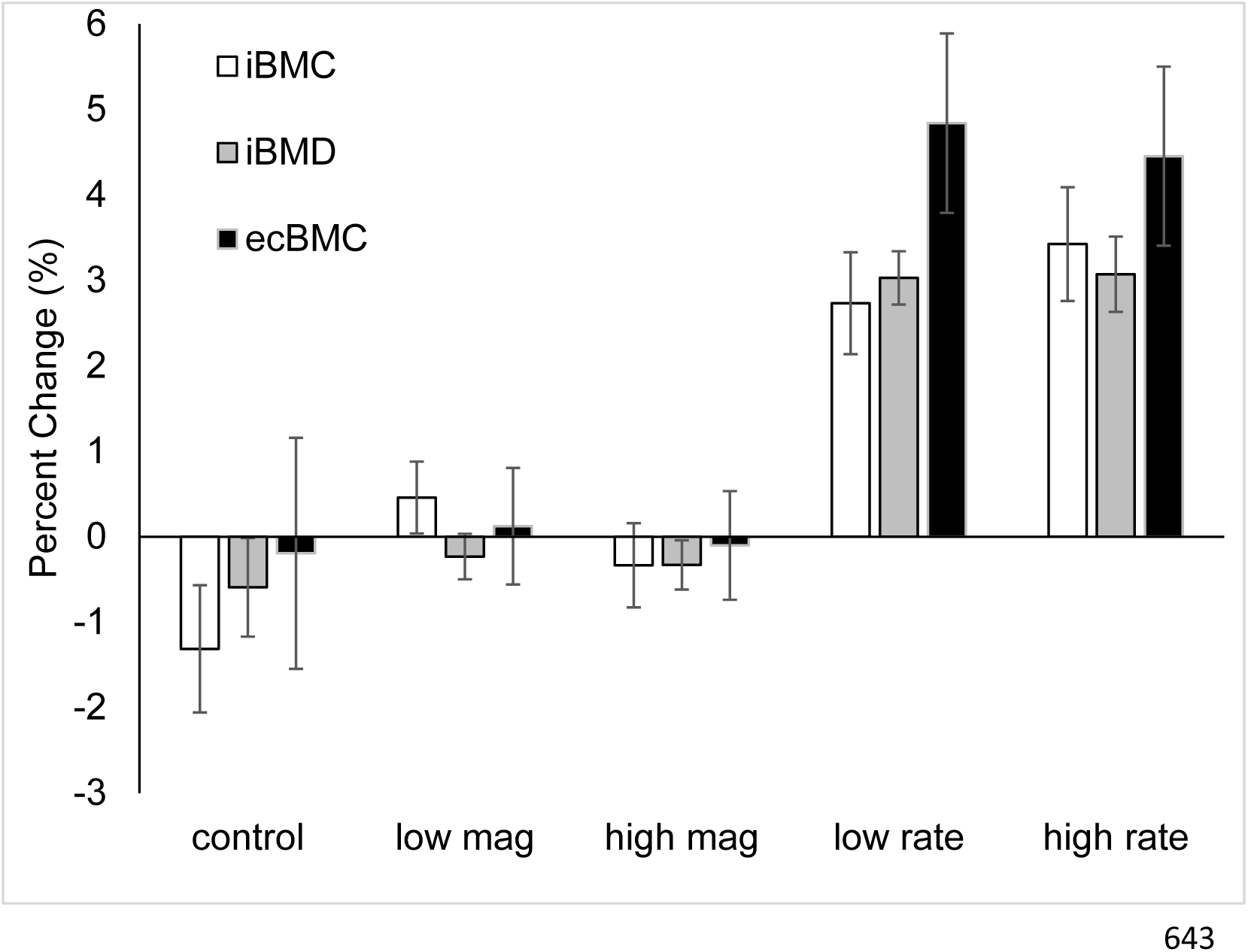
12-month changes in QCT-derived primary outcome variables. Both the low and high rate groups had significant differences compared to the control group in all three variables.

**Table 2.** Mean (SD) baseline of the pooled data, and percent change at 12 months in QCT variables, by group.

|  | Total<br>(n=66)<br>Baseline |  | Control<br>(n=13)<br>%ΔV5 |  | Low Magnitude<br>(n=13)<br>%ΔV5 |  | High Magnitude<br>(n=17)<br>%ΔV5 |  | Low Rate<br>(n=12)<br>%ΔV5 |  | High Rate<br>(n=11)<br>%ΔV5 |  |
| --- | --- | --- | --- | --- | --- | --- | --- | --- | --- | --- | --- | --- |
| Ultradistal |  |  |  |  |  |  |  |  |  |  |  |  |
| iBV (cm³) | 3.84 | (0.40) | -1.17 | (2.51) | 0.75 | (2.14) | -0.01 | (2.23) | -0.29 | (1.29) | 0.35 | (1.86) |
| iBMC (g) | 0.91 | (0.15) | -1.31 | (2.68) | 0.46 | (1.52) | -0.33 | (2.03) | 2.73 | (2.07) | 3.42 | (2.21) |
| iBMD (g/cm³) | 0.24 | (0.03) | -0.59 | (2.08) | -0.23 | (0.96) | -0.33 | (1.19) | 3.03 | (1.08) | 3.07 | (1.46) |
| ecBV (cm³) | 1.99 | (0.15) | -0.41 | (1.49) | 0.89 | (1.39) | 0.34 | (1.58) | -0.01 | (1.25) | 0.41 | (1.46) |
| ecBMC (g) | 0.63 | (0.18) | -0.19 | (4.87) | 0.12 | (2.46) | -0.10 | (2.62) | 4.84 | (3.64) | 4.45 | (3.47) |
| ecBMD (g/cm³) | 0.31 | (0.08) | 0.21 | (4.52) | -0.74 | (2.79) | -0.43 | (2.59) | 4.85 | (3.46) | 4.03 | (3.30) |
| Total |  |  |  |  |  |  |  |  |  |  |  |  |
| iBV (cm³) | 12.96 | (1.55) | -0.06 | (0.47) | -0.25 | (0.52) | -0.13 | (0.45) | 0.16 | (0.64) | 0.38 | (0.41) |
| iBMC (g) | 5.10 | (0.60) | -0.23 | (1.20) | -0.45 | (0.80) | -0.19 | (0.78) | 1.97 | (0.86) | 1.93 | (0.75) |
| iBMD (g/cm³) | 0.39 | (0.04) | -0.17 | (1.14) | -0.19 | (0.58) | -0.06 | (0.78) | 1.81 | (0.63) | 1.55 | (0.86) |
| ecBV (cm³) | 8.24 | (0.72) | 0.17 | (0.71) | -0.07 | (0.57) | 0.18 | (0.44) | 0.45 | (0.62) | 0.35 | (0.64) |
| ecBMC (g) | 4.09 | (0.55) | 0.15 | (1.78) | -0.26 | (1.11) | 0.00 | (1.01) | 2.06 | (1.40) | 1.72 | (1.37) |
| ecBMD (g/cm³) | 0.50 | (0.05) | -0.02 | (1.28) | -0.20 | (0.76) | -0.18 | (0.98) | 1.60 | (0.84) | 1.36 | (0.83) |
| Ultradistal Strength |  |  |  |  |  |  |  |  |  |  |  |  |
| CSA (cm²) | 4.15 | (0.42) | -0.67 | (1.67) | 0.65 | (1.45) | 0.00 | (1.48) | -0.26 | (1.23) | 0.35 | (1.75) |
| CSI (g²/cm⁴) | 0.24 | (0.07) | -1.81 | (4.40) | 0.18 | (1.96) | -0.65 | (2.85) | 5.90 | (3.05) | 6.62 | (3.26) |
| BSI (cm³) | 0.12 | (0.03) | -0.59 | (4.31) | 0.32 | (1.90) | -0.29 | (2.30) | 4.74 | (2.44) | 4.81 | (2.70) |

Data presented as mean (SD). Bold indicates significant regression coefficient representing contrast with control group for raw change
Baseline is for subjects with follow-up data available

**Table 3.** Standardized regression coefficients for QCT, by group and by loading dose. Low1 and High1 indicate low and high strain magnitude groups from Experiment 1. Low2 and High2 indicate low and high strain rate groups from Experiment 2.

|  | **Model Definition** | **R<sup>2</sup>** | **β<sub>1</sub>** | **β<sub>2</sub>** |
| --- | --- | --- | --- | --- |
| **UD iBMC (g)** | β<sub>1</sub>*Low1+β<sub>2</sub>*High1+ε | 0.101 | **0.374** | 0.221 |
|  | β<sub>1</sub>*Low2+β<sub>2</sub>*High2+ε | **0.438** | **0.599** | **0.678** |
|  | β<sub>1</sub>*StrainStim+ε | 0.039 | 0.197 |  |
|  | β<sub>1</sub>*Strain_MagRate+ε | **0.117** | **0.342** |  |
|  | β<sub>1</sub>*Strain_Mag+ε | **0.124** | **0.352** |  |
|  | β<sub>1</sub>*Strain_Rate+ε | **0.102** | **0.320** |  |
| **UD iBMD (g/cm<sup>3</sup>)** | β<sub>1</sub>*Low1+β<sub>2</sub>*High1+ε | 0.009 | 0.091 | 0.101 |
|  | β<sub>1</sub>*Low2+β<sub>2</sub>*High2+ε | **0.563** | **0.739** | **0.716** |
|  | β<sub>1</sub>*StrainStim+ε | 0.003 | 0.051 |  |
|  | β<sub>1</sub>*Strain_MagRate+ε | 0.041 | 0.203 |  |
|  | β<sub>1</sub>*Strain_Mag+ε | 0.045 | 0.212 |  |
|  | β<sub>1</sub>*Strain_Rate+ε | 0.044 | 0.209 |  |
| **Total iBMC (g)** | β<sub>1</sub>*Low1+β<sub>2</sub>*High1+ε | 0.019 | -0.119 | 0.028 |
|  | β<sub>1</sub>*Low2+β<sub>2</sub>*High2+ε | **0.545** | **0.743** | **0.687** |
|  | β<sub>1</sub>*StrainStim+ε | 0.001 | -0.028 |  |
|  | β<sub>1</sub>*Strain_MagRate+ε | 0.008 | 0.091 |  |
|  | β<sub>1</sub>*Strain_Mag+ε | 0.011 | 0.106 |  |
|  | β<sub>1</sub>*Strain_Rate+ε | 0.012 | 0.109 |  |
| **Total iBMD (g/cm<sup>3</sup>)** | β<sub>1</sub>*Low1+β<sub>2</sub>*High1+ε | 0.005 | -0.018 | 0.059 |
|  | β<sub>1</sub>*Low2+β<sub>2</sub>*High2+ε | **0.520** | **0.756** | **0.627** |
|  | β<sub>1</sub>*StrainStim+ε | 0.001 | 0.027 |  |
|  | β<sub>1</sub>*Strain_MagRate+ε | 0.023 | 0.153 |  |
|  | β<sub>1</sub>*Strain_Mag+ε | 0.036 | 0.189 |  |
|  | β<sub>1</sub>*Strain_Rate+ε | 0.012 | 0.166 |  |

Bold For R<sup>2</sup> indicates p<0.05 for F-test of overall model fit
Bold For β<sub>1</sub> or β<sub>2</sub> indicates p<0.05 for t-test of significance for coefficient

In models examining the effects of loading dose on the changes to bone, ultradistal iBMC, iBV, and ecBV were all positively and consistently associated with measures of loading dose, especially Strain_MagRate (Figure 4a). However, in all cases, loading dose explained less than 15% of the variance in the change values.

**Figure 4.**
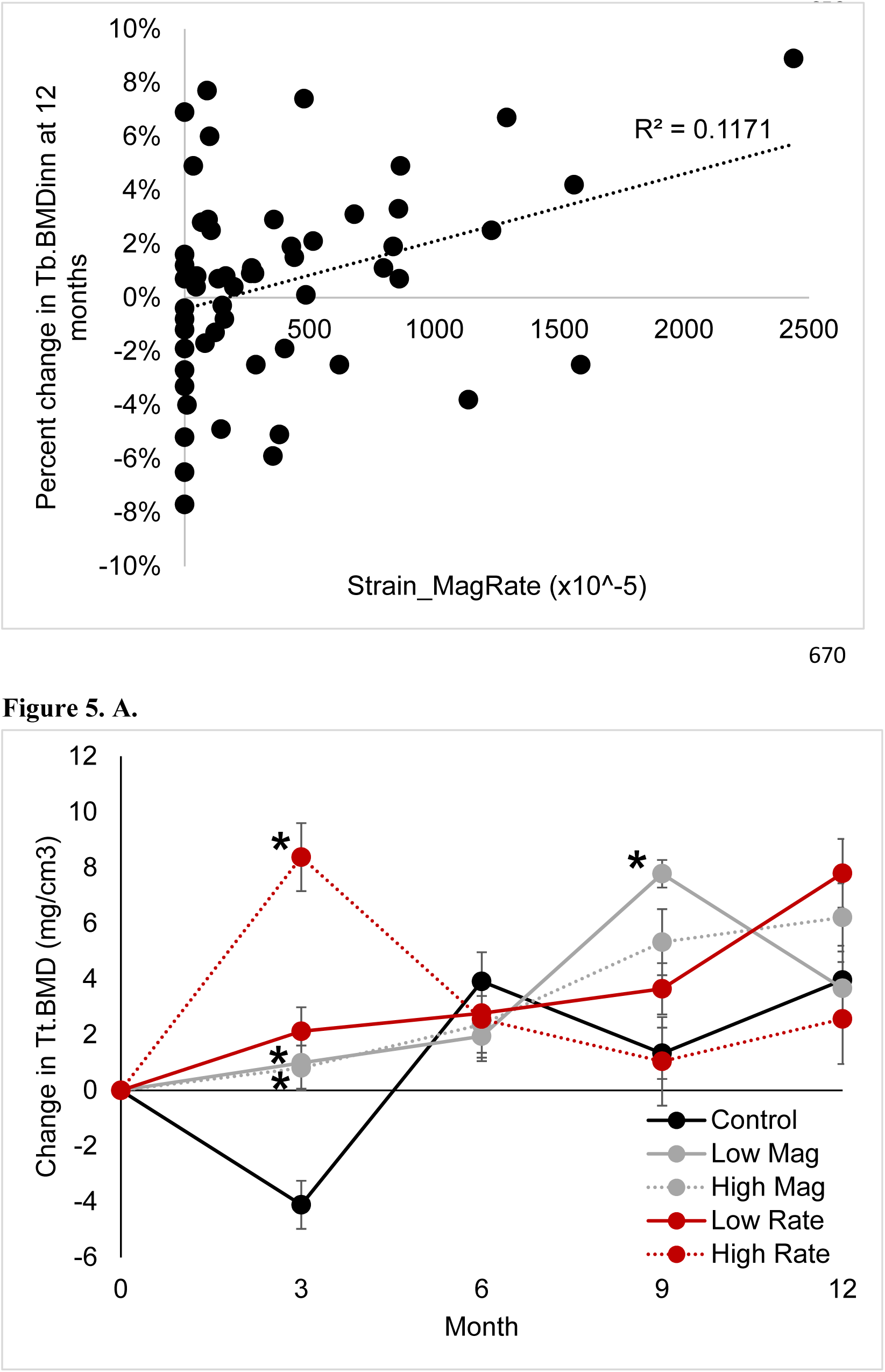
A. Percent change in ultradistal iBMC versus Strain_MagRate. **B.** Percent change in inner trabecular BMD versus Strain_MagRate. Both plots represent 12-month change for all subjects with available data.

### Effect of Strain on 3, 6, 9, and 12 Month Bone Microstructure (HRpQCT)

After three months, membership in the low and high magnitude loading groups explained up to 17% of the increases in Tt.BMD compared to the control group (Supplemental Table 1). Similarly, high loading rate was significantly associated with three-month increases to Tt.BMD, Ct.BMD and Ct.Th (Supplemental Table 1). Strain_MagRate, Strain_Mag, and Strain_Rate were all significant predictors for the change in Tt.BMD and Ct.Th, although 12% or less of the variance in these measures was explained by loading dose.

At six months, none of the microstructural changes were different between groups. However, at nine months, the low strain magnitude group was significantly and positively associated with increases to Tt.BMD, Tb.BMD, Tb.BMDinn, and Tb.BMDmeta (Figure 5). Similarly, the high strain magnitude and low strain rate groups were positively associated with changes to Tb.BMDinn. Strain_MagRate and Strain_Rate, and were also positively associated with the increase to Tb.BMDinn in nine months. These changes persisted at 12 months, with Strain_MagRate being associated with increases to Tb.BMD and Tb.BMDinn (Figure 4b).

**Figure 5.**
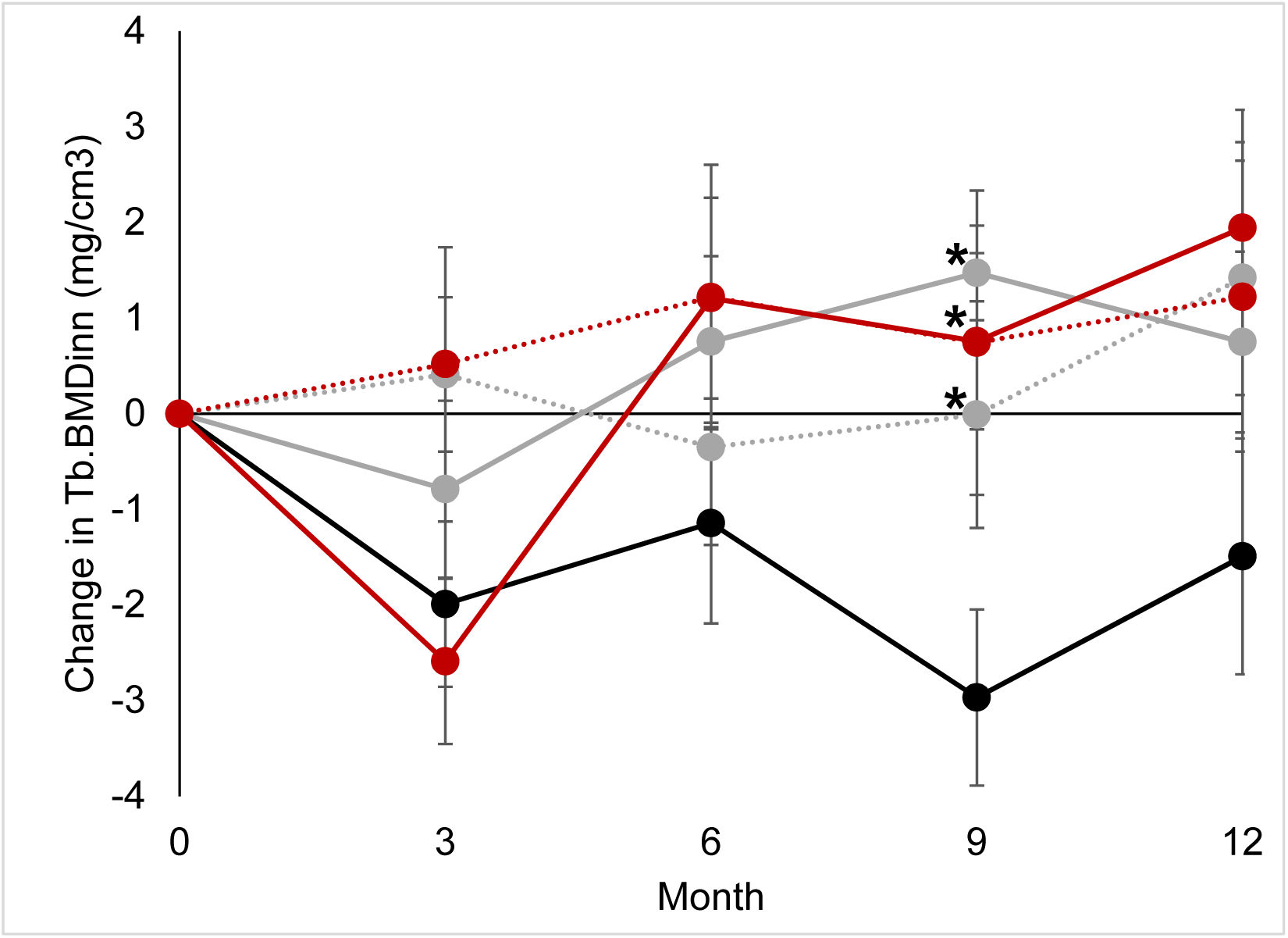
A. Change in Tt.BMD, and **B.** Tb.vBMDinn from baseline versus time, per group. Significant group changes versus control group at specific timepoints are labeled with *. Error bars represent standard error.

### Comparison between change in ultradistal iBMC tertile groups

Subjects in the first tertile gained bone, the middle tertile had no change, and the lowest tertile lost bone. Subject age, height, weight, aBMD at baseline, and vitamin D levels were not different between tertile groups (Table 4). Those in the first tertile had higher baseline ultradistal iBMC and iBMD than the lowest tertile, but no other baseline measures differed between groups. All loading-related variables except StrainStim were significantly different across the three tertiles, with the first tertile having the highest dose. However, after Bonferroni adjustment for multiple comparisons, only the first and third tertiles showed significant differences (Figure 6).

**Figure 6.**
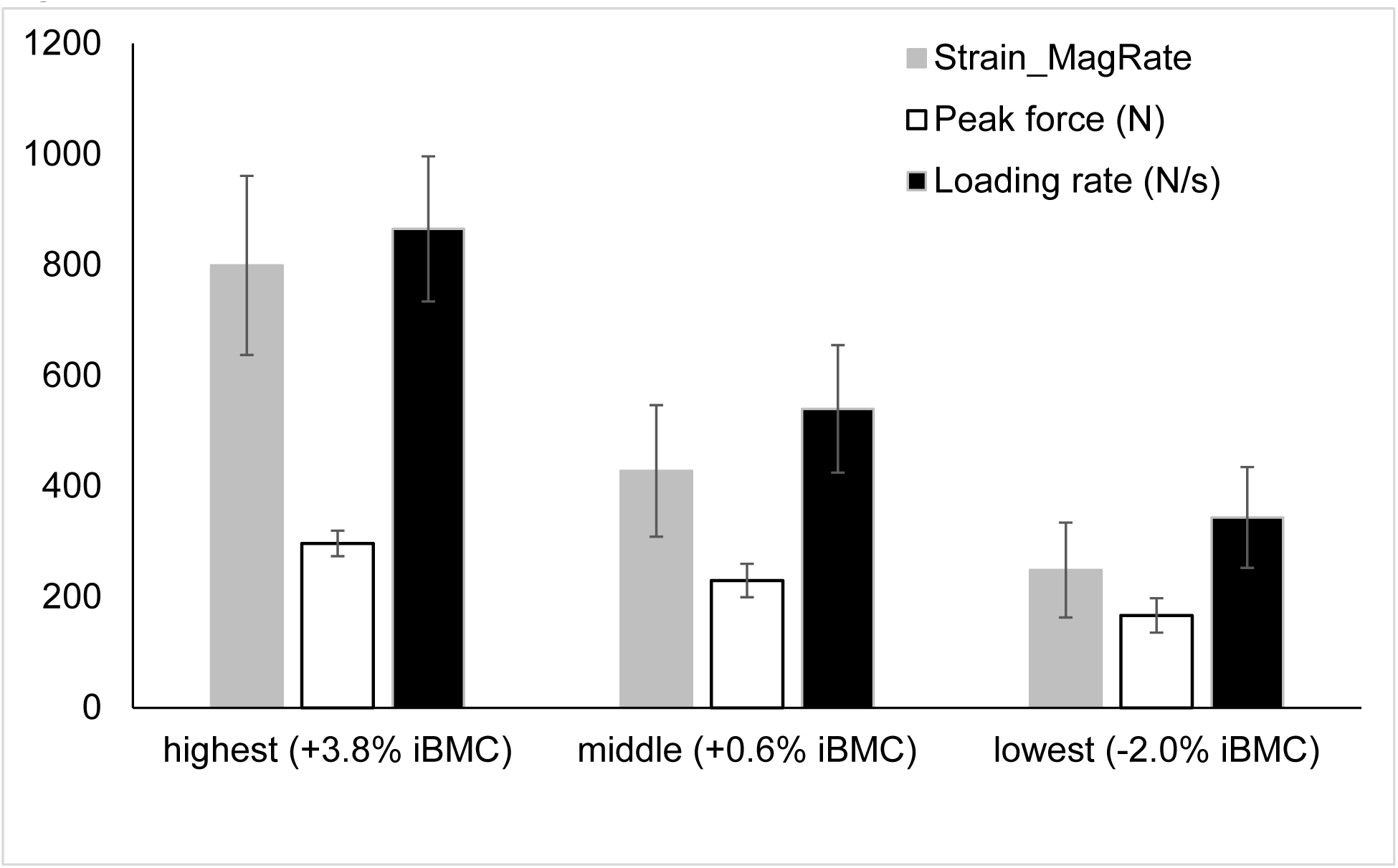
Comparison of key loading metrics between the first, middle, and lowest gain in iBMC tertile. For all variables shown, the first and lowest tertiles were significantly different from each other after Bonferroni correction.

**Table 4.**
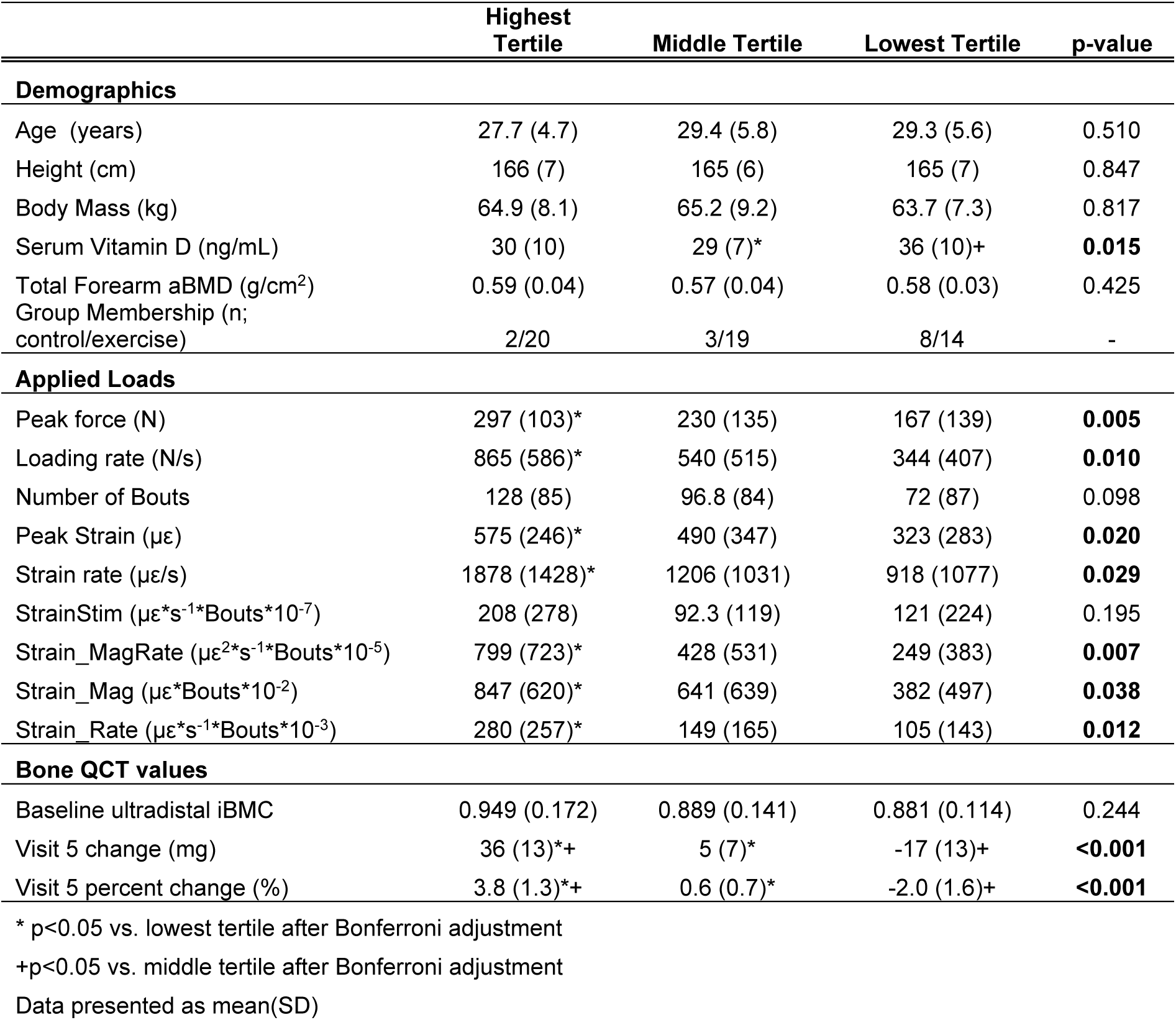
Mean (SD) grouped by change in ultradistal iBMC tertile. P-values indicate significant between-group differences. Symbols indicate significant Bonferroni-adjusted post hoc comparisons between specific tertiles.

## Discussion

We conducted a randomized prospective trial to characterize the relationship between mechanical strain magnitude and rate and changes to bone in healthy adult women. We found that the application of mechanical strain produced small but significant changes to the ultradistal radius after one year. Our first hypothesis was partially supported by the results. Membership in both the low and high strain rate groups were strongly associated with increases in bone mass and density. Both low and high strain rates were also associated with increases to compressive and bending strength indices. When loading dose, which included a combination of strain magnitude, strain rate, and number of loading bouts was taken into account, we observed a dose-dependent relationship between iBMC and CSA across all subjects.

Our hypothesis that structural changes would include increased cortical diameter and thickness, and increased trabecular bone mass near the endosteal surface was only partly supported. Endocortical BV, BMC, and BMD increased at 12 months, indicating bone apposition on both the periosteal and endosteal surfaces due to loading. At 3, 9, and 12 months, increases to overall density and trabecular density were observed with HRpQCT, and were dependent on loading dose. However, contrary to our expectation, the inner trabecular density (Tb.BMDinn) rather than more peripheral regions appeared to be primarily affected. During aging, trabecular structure is first lost from this region, and later from more peripheral regions^(37)^, thus maximizing moment of inertia for a given quantity of bone. We previously reported age-associated declines in Tb.BMDinn within a large subset of the subjects measured here ^(38)^. It is possible that in our cohort of young, healthy women, trabecular microstructure in the more peripheral regions was already at its physiologic maximum, limiting the degree to which it might be improved. However, even in this group, we observed Tb.BMDinn was lower than Tb.BMDmeta (Supplemental Table 1), suggesting that there was greater capacity to improve the inner region with anabolic physical activity.

We observed significant positive effects of loading on Tt.BMD, Tb.BMDinn, and Ct.BMD after three months. Interestingly, all subjects were assigned the same loading magnitude (200 N) during this ramp-up period, rather than a group-specific strain. Compliance was also the best during the first three months. Therefore, it is not surprising that both low and high magnitude groups had increases in to these variables, since they both received the same stimulus. Overall, this supports the notion that loads must be novel to elicit an osteogenic response ^(18)^. The improved response in the low magnitude group, who completed more loading bouts than other groups, also suggests that consistency of exercise is as important as strain magnitude and rate.

We observed significant increases to bone mass in the strain rate experiment; however, both low and high strain rate groups demonstrated positive results and the regression coefficients were similar between groups. Surprisingly, in Experiment 1 (strain magnitude) only the low strain magnitude group showed even slight increases in ultradistal iBMC after 12 months, with no observable changes in the high strain magnitude group. In fact, despite being given different target strains and strain rates, the different loading groups did not achieve the expected range of rates and magnitudes (Table 1). This, combined with varying subject compliance may partly explain these counterintuitive results. The analysis by tertile change in ultradistal iBMC suggests that, despite variable results, changes to bone are indeed associated with bone loading dose. Subjects in the highest tertile also had had a non-significant trend towards higher baseline BMC, suggesting that perhaps these individuals simply had a greater physiologic capacity to respond to osteogenic stimuli. We did not observe any other obvious factors related to the change (e.g. vitamin D status) that might explain this, although our measurements did not include biomarkers related to bone metabolism. The degree to which strain magnitude can be manipulated is limited due to risk of secondary injury, although greater magnitudes are possible in the lower extremities. With vibration and other external assistance, it is possible to manipulate strain rate over a much wider range than strain magnitude.

In contrast to small animal *in vivo* loading models, which use a materials testing machine to generate a predictable and repeatable waveform, voluntarily applied forces are variable in terms of frequency content, even when the peak magnitude is guided through visual feedback, as in our study. While many measures of bone loading dose have been proposed in the literature^(18, 21, 22, 36)^, we found it impractical to implement any of them exactly as described by the authors. In addition to voluntarily produced loading signals being inconsistent, mechanical strain is non-uniform within a bone, both temporally and spatially; thus, no single strain value completely describes the strain occurring within a bone. Furthermore, it is not practical to place strain gages on most bones, and even when such measures are obtained (e.g. reference ^(39)^), they only represent a small fraction of the bone surface. Here, we examined several candidate versions of loading dose, based on recorded load cell signals and subject-specific FE models. Each version included a combination of strain magnitude, frequency, and number of loading bouts. As a first attempt, we chose to examine the continuum strain produced within the analysis region in question (corresponding with the QCT or HRpQCT analysis region for those respective variables) and the total number of bouts achieved up to the timepoint in question. While we found significant associations between our measures of loading dose and changes to bone in our subjects, we found that at best, dose explained 12% of the variance in the change. It is possible that other formulations of loading dose that include local strain rate, strain gradient, or other measures, may be more relevant.

The magnitude and nature of the changes we observed are similar to an earlier, 6-month study using a similar loading protocol ^(26)^. In that set of 19 young women, control subjects lost 1.7±1.1% ultradistal iBMC, while those in the loading group had no change in iBMC, but significant increases in trabecular BMC (1.3±2.8%). Here, we found a similar decrease in the control group iBMC (-1.3±2.7%), and increases to BMC and BMD that were associated with loading dose. The present cohort differed from the previous study in several ways. First, present subjects were generally older (28 vs. 22 years old) and many had a history of pregnancy or lactation (although not within the two years preceding enrollment). The present group were assigned loading magnitudes based on strain within the ultradistal radius at the instant of peak force production. However, due to limits on the force that subjects were able to safely produce voluntarily, subjects fell short of their target strains. Thus, while high strain magnitudes may have, in theory, elicited a greater osteogenic response, they were impractical or unsafe to implement. Similarly, subjects in the low and high strain rate groups were given instruction sets designed to elicit significantly different strain rates. While the rates were significantly different between groups, (low: 675 με/s, high: 1430 με/s) the sample did not vary as widely as designed. And, both low and high strain rate groups experienced similar increases in bone. Distal radius compressive loading is a relatively constrained activity, and the ability to manipulate the strain signal was limited.

Our results suggest that, while compressive loading in general is osteogenic, it is not necessary to generate extremely high strain magnitudes or rates to elicit a positive response in the upper extremity. Significant gains in BMC were associated with moderate strain rates and magnitudes and can be achieved in a reasonable amount of time (100 loading cycles/bout, and an average of 131 loading bouts over a 12-month period for the highest tertile group). This is reassuring, since high loading rates have been linked to increased risk of stress fracture ^(40)^. Although we did not systematically test the effect of loading cycles/bout, we based our target of 100 on (1) feasibility and time to complete the intervention, about three minutes, and (2) theoretical calculations of bone adaptation ^(36, 41)^ that suggested a diminished osteogenic response with additional cycles. It is possible that the *number* of repetitive loading cycles, more than the strain signals from individual loading cycles themselves, is what tips the balance of a strain signal being positive/osteogenic to negative/increasing risk of injury.

This study had several important limitations. Only 66 of the 102 original subjects completed all 12 months of the study, and the results may be biased towards those who did not drop out. However, the demographics and baseline data of individuals who dropped out were not different from those who completed the study. Due to the lower number of completers, we were not powered to detect trabecular microstructural changes. However, we did observe significant changes to iBMC and Tb.BMD. The magnitude of the increases to iBMC due to the loading interventions, 1.2% across all subjects, is not dissimilar to other treatment effects considered clinically relevant. And, subjects participating in Experiment 2 had much larger increases (2.9% and 3.6%). For comparison, a 3.3% increase in trochanter integral BMD over 36 months was observed in postmenopausal women given zolendronic acid ^(42)^, and it has been estimated that each 1% increase in peak bone mass imparts over 1 year of osteoporosis-free life in the future^(43)^. While the present study examined the effects of strain magnitude and rate on bone adaptation, an underlying assumption is that the bone of each individual is already well adapted for her habitual activities; our analysis only considered the novel/added stimulus. Although we collected physical activity data as part of this study, they were beyond the scope of the present analysis, but may potentially explain some of the variability in response to our intervention. Our results may not be generalizable to other populations, including postmenopausal women, those with low vitamin D, men, or specific clinical populations. Finally, more research is needed to determine the specific strain requirements to elicit clinically relevant changes to lower extremity bone, given the high habitual loading stimulus in these anatomic sites.

Although other clinical trials have investigated the efficacy of various types of exercise to for improving bone mass, this study is the first to systematically investigate the effect of mechanical strain rate and magnitude on bone adaptation in humans. The data presented here fill a critical translational gap, linking *in vivo* animal models to clinical trials, and may be useful for informing the design of future clinical interventions for bone health. In particular, our data show that in healthy adult women, the distal radius is capable of modest adaptation in response to mechanical strain, and that the adaptation is associated with measures of loading dose that include strain magnitude, rate, and number of loading bouts.

In conclusion, we conducted a randomized prospective experiment to systematically investigate the effect of mechanical strain rate and magnitude on bone adaptation, using an *in vivo* upper extremity loading model in healthy adult women. We found that compressive loading was osteogenic, with high and low strain rate groups having similar significant increases to bone mass. We observed that subjects who gained the most bone had, on average, completed 130 compressive loading bouts, generating an average energy-equivalent strain of 550 με at 1805 με/s within the distal radius, over a period of 12 months. Individuals with the greatest gains to bone mass were similar in demographics to those with the lowest gains to bone mass. We conclude that signals related to strain magnitude, strain rate, and number of loading bouts collectively contribute to bone adaptation in healthy adult women.

---

**Supplemental Table 1.**
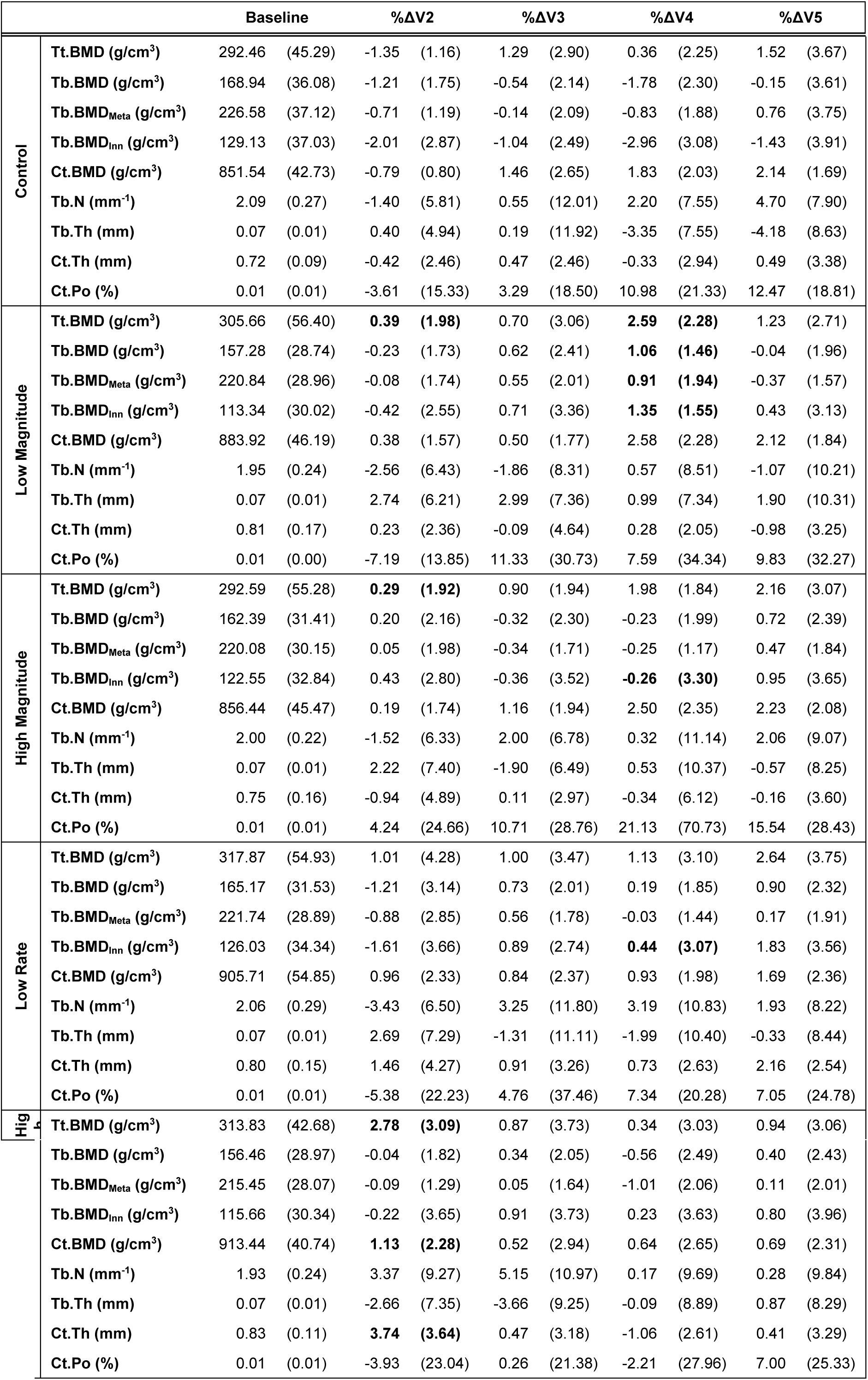
Mean (SD) of baseline and percent changes of HRpQCT measures during each visit (V1 at baseline through V5 at 12 months), by group.

## Acknowledgements

This research was fully supported by NIAMS of the National Institutes of Health under award number R01AR063691.The content is solely the responsibility of the authors and does not necessarily represent the official views of the National Institutes of Health. We thank Sabahat Ahmed for her organization and dedication as research coordinator, and Dr. Jane Marone for serving as our Independent Safety Monitor.

## Author Contributions

Study conceived by KLT and designed by KLT with assistance from TJS. Data collection by MEM, KLT, JEJ, and TAB. Data analysis and interpretation: MEM, KLT, JEJ, TAB, ZW, TJS. Manuscript writing: KLT and MEM. Manuscript approval: MEM, KLT, JEJ, TAB, ZW, TJS.

